# RiboPipe: efficient per-transcript codon-resolution ribo-seq coverage imputation for low-coverage transcripts

**DOI:** 10.64898/2026.03.20.711481

**Authors:** Yao-zhong Zhang, Satoshi Hashimoto, Sihan Li, Toshifumi Inada, Seiya Imoto

## Abstract

**Motivation:** Ribosome profiling (Ribo-seq) provides codon-resolution measurements of translation; however, many transcripts exhibit sparse or low read coverage, which limits downstream quantitative analyses. Reliable prediction and imputation of codon-resolution coverage for low-coverage transcripts remain computationally challenging.

**Results:** We present RiboPipe, an efficient framework for per-transcript codon-resolution Ribo-seq coverage imputation for low-coverage transcripts. RiboPipe is designed around three key principles. First, it jointly optimizes transcript-level mean ribosome load (MRL) prediction and codon-level coverage modeling within a unified objective, enabling consistent learning across both local and transcript-level scales. Second, it introduces a peak-weighted loss that emphasizes high-signal codon positions associated with translational pausing, improving the recovery of functionally relevant coverage peaks. Third, the framework is lightweight and data-efficient, achieving stable performance even when trained on only a small fraction of high-coverage transcripts. Based on two publicly available Ribo-seq datasets (GSE233886 and GSE133393) as instances, we demonstrate stable convergence and consistent prediction accuracy across multiple train–test split ratios. Comparative evaluation of embedding strategies shows that simple one-hot representations achieve superior performance compared with pre-trained language model embeddings under identical training conditions. Overall, RiboPipe provides a computationally efficient and scalable framework for Ribo-seq coverage imputation in low-coverage transcripts.

**Availability and Implementation:** The source code and associated data can be accessed at https://github.com/yaozhong/riboPipe

## 1 Introduction

Ribosome profiling (Ribo-seq) enables measurement of translation by sequencing ribosome-protected fragments, providing ribosome density profiles with codon-level resolution. Codon-resolved footprints have been widely used to study translation elongation dynamics, including pausing events and ribosome collisions that may arise from congestion on transcripts. However, in typical Ribo-seq experiments, many transcripts exhibit sparse or low ribosome footprint coverage due to low transcript abundance, limited sequencing depth, uneven library complexity, and condition-dependent translation. This sparsity is particularly problematic when the analysis requires accurate reconstruction of local high-signal positions (“peaks”) that are informative for elongation speed and potential collision propensity.

A broad ecosystem of computational tools supports Ribo-seq processing and downstream inference, including methods for A-site assignment and translation efficiency estimation that account for sampling errors and biases (e.g., Scikit-ribo) (Fang *et al*., 2018). Other probabilistic approaches focus on detecting translated ORFs by modeling positional coverage patterns (Choudhary *et al*., 2020). While these methods are essential for standard analysis, they are not designed to directly model or reconstruct transcript-wide ribosome occupancy at codon resolution, particularly under sparse coverage conditions.

Recently, deep learning has been applied to model translation from sequence. For transcript-level translation outputs, mean ribosome load (MRL) has been predicted from regulatory sequence contexts such as 5’ UTRs (Karollus *et al*., 2021). For codon-resolution density modeling, several models predict full-length ribosome density profiles across CDS regions. RiboMIMO introduced a multi-input, multi-output neural architecture that predicts ribosome density distributions across codons within a CDS sequence (Tian *et al*., 2021). More recently, transformer-based approaches such as Riboformer demonstrated strong performance in predicting codon-level ribosome densities and highlighted links between learned signals and elongation-related phenomena, including collision-associated determinants (Shao *et al*., 2024). Hybrid frameworks also explore the integration of mechanistic simulation with machine learning for generating ribosome count profiles (e.g., seq2ribo) (Kaynar and Kingsford, 2026). These studies suggest that codon-level ribosome density can be largely explained from sequence features and regulatory context under well-covered conditions.

Nevertheless, several practical challenges remain for routine use with typical Ribo-seq datasets. First, many applications require reliable inference for *low-coverage transcripts*, where training signals are limited and naive models tend to under-recover sharp peaks that may correspond to slow codons or biologically meaningful pauses. Second, existing modeling efforts often optimize profile prediction alone, without explicitly coupling codon-level profile recovery to a transcript-level translation summary such as MRL, which can provide a stable supervisory signal under sparse observations. In addition, for practical deployment as a software tool, there is a need for lightweight and data-efficient training and inference strategies that perform well with limited training data and standard computational resources.

To address these needs, we present **RiboPipe**, an efficient framework for per-transcript codon-resolution Ribo-seq coverage imputation of low-coverage transcripts. RiboPipe is designed around three principles. (i) *Joint optimization across scales*: the model simultaneously learns transcript-level MRL prediction and codon-level coverage modeling within a unified objective, improving stability under sparse observations. (ii) *Peak-weighted optimization for elongation dynamics*: a peak-weighted loss emphasizes high-signal codon positions that are most informative for translation speed and related elongation phenomena, improving recovery of functionally relevant peaks. (iii) Lightweight and data-efficient training: the framework is engineered to achieve stable performance even when trained on a small fraction of high-coverage transcripts, enabling practical use on typical datasets. In addition, we provide a controlled comparison of embedding strategies (one-hot versus pre-trained language model representations) under identical training protocols to inform practical deployment choices.

## 2 Methods

### 2.1 Problem setting and application scenario

RiboPipe is designed for within-sample imputation of codon-resolution ribosome coverage. Given a single Ribo-seq sample, transcripts with high coverage provide reliable codon-level ribosome occupancy profiles, whereas many transcripts exhibit sparse or low coverage due to limited sequencing depth. Under the same cellular condition, global translational determinants such as tRNA abundance, elongation factor availability, and ribosome pool composition are shared across transcripts. Consequently, codon usage patterns and elongation-related sequence determinants learned from high-coverage transcripts can be transferred to predict codon-level coverage in low-coverage transcripts. Formally, let *𝒯*_high_ denote transcripts with sufficient coverage and *𝒯*_low_ denote low-coverage transcripts within the same sample. The objective is to learn a mapping from coding sequence to codon-resolution ribosome coverage using *𝒯*_high_, and then apply the learned model to infer coverage for *𝒯*_low_.

### 2.2 Coverage normalization and target definition

For each transcript *t*, raw Ribo-seq read counts were obtained at codon resolution. To focus on relative ribosome occupancy patterns rather than absolute expression levels, counts were normalized within each transcript:

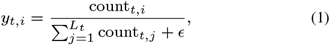

where *L*_*t*_ denotes the CDS length in codons and ∈ prevents division by zero. This normalization encourages the model to learn codon-level occupancy distributions independent of transcript abundance. The transcript-level mean ribosome load (MRL) is defined as: 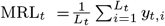. RiboPipe jointly predicts both codon-resolution coverage ŷ_*t*_ and transcript-level mean ribosome load 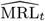, enabling simultaneous modeling of local occupancy patterns and global translational activity.

### 2.3 Sequence representation and biological features

Each codon in the CDS is represented either as a one-hot encoding or as a context-aware embedding derived from a pre-trained codon language model. In addition to nucleotide sequence features, we incorporate biologically motivated features, following previous work Aguilar Rangel *et al*. (2024). These features are concatenated with the sequence embeddings and include codon frequency, the tRNA adaptation index (tAI), wobble decoding indicators, and amino acid physicochemical properties such as hydrophobicity, polarity, and charge.

### 2.4 Model architecture and lightweight design

RiboPipe adopts a compact bidirectional LSTM backbone to model contextual dependencies along the coding sequence. Codon embeddings are processed sequentially, and two regression heads are applied: one predicts codon-level normalized coverage, and the other predicts transcript-level MRL through sequence pooling. The architecture is intentionally lightweight to enable stable training with a limited number of high-coverage transcripts, without requiring large-scale pretraining.

### 2.5 Joint peak-weighted optimization

Training is performed using a joint objective: ℒ = ℒ_cov_ + ℒ_MRL_, where the transcript-level loss is defined as:

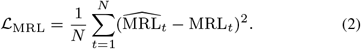

To improve recovery of high-signal codon positions associated with elongation slowdown or ribosome pausing, a peak-weighted mean squared error is used for codon-level prediction:

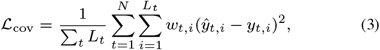

with weights defined as: 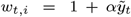, *i*, where 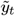,*i* denotes the normalized coverage magnitude. This peak-weighted objective improves recovery of functionally relevant high-occupancy codons while maintaining stability in low-signal regions.

## 3 Results

### 3.1 Datasets and experimental setup

Experiments were conducted using publicly available Ribo-seq datasets generated under controlled cellular conditions. We evaluated our framework on GSE233886 Schneider-Poetsch *et al*. (2025) and GSE133393 Han *et al*. (2020), derived from HEK293F and HEK293T cell lines, respectively. GSE233886 exhibits strong three-nucleotide periodicity and relatively smooth elongation signals, whereas GSE133393 shows higher-variance coverage patterns associated with ribosome collisions. Due to page limitations, results for GSE133393 are presented in the Supplementary Materials. Raw ribosome footprint reads were aligned to coding sequences annotated in GENCODE v47, followed by P-site assignment to obtain codon-resolution ribosome occupancy profiles. P-site positions and counts were extracted using riboWaltz Lauria *et al*. (2018). Transcripts were ranked by total CDS coverage; those above the 75th percentile (P75) were used for training, while the remainder were reserved for evaluation. Within the high-coverage set, transcripts were further split into training and validation subsets (e.g., 80/20). Model performance was assessed using mean squared error (MSE) and Pearson correlation at both codon and transcript levels. To evaluate biologically relevant high-signal regions, we additionally computed peak-oriented metrics on the top 5% highest-occupancy codons.

### 3.2 Codon-resolution prediction performance

Under the evaluation protocol described above, RiboPipe demonstrated stable optimization behavior and consistent codon-resolution reconstruction performance (Figure 1a). As shown in Fig. 1 for GSE233886, across 200 training epochs both standard MSE and peak-weighted MSE decreased smoothly without oscillatory instability. In parallel, binary classification accuracy increased steadily and reached saturation after approximately 150 epochs. At the transcript level, MRL prediction exhibited rapid early-phase improvement followed by gradual refinement. Both Pearson and Spearman correlations increased sharply within the first 50 epochs and stabilized thereafter, demonstrating effective capture of global translational trends in addition to local codon-level structure. Case studies from the test set (shown in Supplementary Fig. S1) demonstrate that the predicted ribosome coverage profiles closely follow the observed codon-resolution signals across transcripts, indicating that the model accurately reconstructs both global coverage patterns and local ribosome occupancy peaks.

**Fig. 1:**
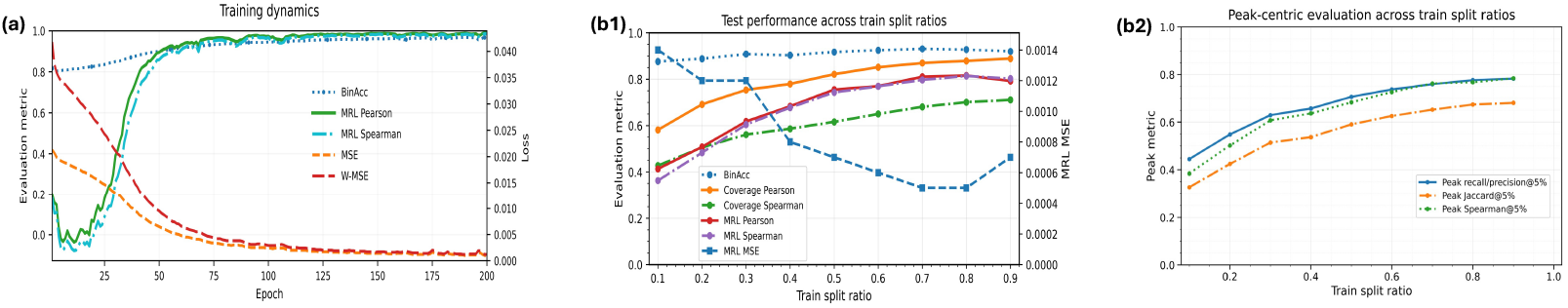
**(a)** Training curves on GSE233886 over 200 epochs, showing binary classification accuracy (BinAcc), transcript-level MRL Pearson and Spearman correlations, and optimization losses (MSE and peak-weighted MSE). **(b1)** Standard test performance across training split ratios, including codon-level coverage Pearson and Spearman correlations, transcript-level MRL correlations, MRL MSE, and binary classification accuracy. **(b2)** Peak-centric evaluation across training split ratios, computed on the top 5% highest-occupancy codons, reporting peak recall and precision overlap, Jaccard similarity, and peak-level Spearman correlation.

### 3.3 Robustness under varying train–test split ratios

We evaluated model robustness across varying train–test split ratios within the high-coverage subset (Figure 1(b1)). Increasing the proportion of training data consistently improved codon-level coverage correlations, with Pearson correlation rising progressively from low-data regimes and approaching near-saturation at intermediate splits. Transcript-level MRL correlations exhibited a similar monotonic trend. Meanwhile, MRL MSE decreased steadily as training fractions increased, indicating improved global regression fidelity. Binary classification accuracy also improved consistently and remained stable across moderate-to-high splits.

To specifically assess recovery of high-occupancy codon positions, we further examined peak-centric metrics defined on the top 5% highest-coverage codons (Figure 1(b2)). Peak overlap (recall/precision) and Jaccard similarity increased monotonically with training fraction, while peak Spearman correlation demonstrated consistent gains across splits. Notably, performance degraded smoothly rather than abruptly under reduced training sizes, suggesting that the joint codon–MRL objective stabilizes learning even in low-data regimes.

### 3.4 Ablation study

We conducted systematic ablations under the default configuration (train split = 0.8; maximum CDS length = 5000 codons) to quantify the contributions of biological features (bioFeat), peak-weighted reconstruction loss (W-MSE), and transcript-level supervision via the MRL head (Table 1). We evaluated five variants: Default, w/o bioFeat, w/o W-MSE (standard MSE only), w/o MRL, and w/o W-MSE+MRL.

**Table 1.** Ablation study on GSE233886 under the default setting (train split = 0.8; max CDS length = 5000 codons). Cov-P/S: codon-level coverage Pearson/Spearman. MRL-P/S/MSE: transcript-level mean ribosome load. PeakOv@5%: overlap between top-5% predicted and true codons. PeakBias@5%: mean(pred–true) on true peak positions (closer to 0 is better).

| Variant | Cov-P | Cov-S | MRL-P | MRL-S | MRL-MSE | BinAcc | PeakOv@5% | PeakJac@5% | PeakSpr@5% | PeakBias@5% |
| --- | --- | --- | --- | --- | --- | --- | --- | --- | --- | --- |
| Default | 0.8788 | 0.7006 | 0.8159 | 0.8136 | 0.0005364 | 0.9274 | 0.7765 | 0.6738 | 0.7674 | -0.0213 |
| w/o bioFeat | 0.8746 | 0.6879 | 0.8035 | 0.8049 | 0.0005456 | 0.9295 | 0.7674 | 0.6605 | 0.7611 | -0.0418 |
| w/o W-MSE | 0.8852 | 0.7348 | 0.8219 | 0.8237 | 0.0004832 | 0.9311 | 0.7704 | 0.6641 | 0.7419 | -0.0471 |
| w/o MRL | 0.8761 | 0.6908 | 0.1453 | 0.1174 | 0.02197 | 0.9263 | 0.7638 | 0.6542 | 0.7554 | -0.0420 |
| w/o W-MSE+MRL | 0.8829 | 0.7364 | 0.5494 | 0.5561 | 0.02492 | 0.9318 | 0.7717 | 0.6665 | 0.7449 | -0.0466 |
| One-hot + bioFeat + UTR | 0.8535 | 0.6358 | 0.7140 | 0.7468 | 0.0008849 | 0.9234 | 0.7394 | 0.6210 | 0.7282 | -0.0811 |
| CodonLM embedding | 0.0312 | 0.0254 | -0.0028 | -0.0071 | 0.0020915 | 0.8348 | 0.0646 | 0.0354 | 0.0100 | -0.4481 |

#### Standard evaluation metrics

Removing W-MSE yielded slightly higher global correlations (Cov-P: 0.8852 vs. 0.8788; Cov-S: 0.7348 vs. 0.7006) and marginally improved MRL correlations (MRL-P: 0.8219 vs. 0.8159), suggesting that standard MSE can optimize overall agreement with the mean coverage profile. In contrast, removing the MRL head substantially impaired transcript-level prediction (MRL-P: 0.1453; MRL-MSE: 0.02197), confirming that explicit transcript-level supervision is essential for capturing global translational trends. Removing bioFeat caused consistent, though modest, decreases in both codon- and transcript-level correlations (Cov-P: 0.8746; MRL-P: 0.8035), indicating complementary predictive information beyond CDS sequence encoding.

#### Peak-centric evaluation

To better reflect peak-sensitive objectives, we assessed peak-oriented metrics on the top 5% highest-occupancy codons. The Default model achieved the best peak recovery and calibration (PeakOv@5%: 0.7765; PeakJac@5%: 0.6738) with the smallest peak underestimation (PeakBias@5%: −0.0213) and the highest peak Spearman correlation (PeakSpr@5%: 0.7674). Notably, w/o W-MSE exhibited stronger global correlations but substantially increased peak shrinkage (PeakBias@5%: −0.0471) and reduced peak rank agreement (PeakSpr@5%: 0.7419), indicating that peak-weighting primarily improves high-occupancy amplitude calibration rather than overall correlation.

### 3.5 Effect of sequence embedding strategies

To evaluate the impact of sequence representation, we compared three encoding strategies: (i) direct codon one-hot encoding, (ii) pretrained codon language model (codonLM) embeddings, and (iii) models incorporating 5^’^UTR features. All experiments used the same LSTM backbone and identical training settings.

#### One-hot codon encoding provides a strong baseline

Using CDS one-hot encoding together with biological features produced stable and accurate codon-level predictions (Pearson ≈ 0.88). Removing biological features led to consistent but modest decreases in performance, indicating that codon identity largely captures the core predictive patterns.

#### Pretrained codon embeddings do not improve performance

Replacing one-hot encoding with pretrained codon language model embeddings substantially degraded predictive accuracy (Pearson ≈ 0.03). One likely explanation is that such embeddings are typically high-dimensional and increase the number of learnable parameters in the downstream model, which can hinder effective learning under the relatively small sample sizes typical of ribosome profiling datasets. In contrast, one-hot encoding directly preserves codon identity and provides a strong inductive bias for learning codon usage patterns within a single cellular condition, which are closely linked to translation elongation dynamics.

#### Limited contribution of 5^’^UTR features for codon-resolution prediction

Incorporating 5^’^UTR sequence features did not improve performance and instead led to consistent decreases in both codon-level coverage and transcript-level metrics. Although 5^’^UTR sequences are known to influence mean ribosome loading through translation initiation, the present task focuses on codon-resolution occupancy along the CDS, which is primarily governed by elongation dynamics. In our experiment, 5^’^UTR information is incorporated as a global transcript-level feature and broadcast across codon positions, which may limit its ability to capture position-specific interactions between initiation context and elongation behavior. Consequently, CDS sequence features remain the dominant determinants of ribosome density patterns, and the additional contribution of 5^’^UTR features is limited under this modeling scheme.

### 3.6 Running time

We evaluated the end-to-end computational cost of RiboPipe using a single ribosome profiling sample (HEK293F WT DMSO) from the GSE233886 dataset under the default configuration (train split = 0.8; maximum CDS length = 5000 codons; 200 training epochs). After applying the CDS length filter (5000 codons), 6,328 transcripts were retained for training. All experiments were conducted on a workstation equipped with an NVIDIA RTX Titan GPU and an Intel Core i9-13900KS CPU.

The complete workflow—including preprocessing, coverage matrix construction, biological feature extraction, and model training—required a total wall-clock time of 911.6 seconds (approximately 15.2 minutes). Neural network training constituted the dominant computational cost (738.9 s; 81.1% of total runtime), whereas preprocessing required 138.8 s (15.2%). Coverage matrix construction (6.9 s) and biological feature extraction (26.9 s) introduced minimal additional overhead.

## 4 Conclusion

In this work, we present RiboPipe, a lightweight framework for within-sample imputation of codon-resolution ribosome occupancy profiles. By learning sequence-dependent translation patterns from high-coverage transcripts, RiboPipe enables accurate and data-efficient reconstruction of codon-level coverage for low-coverage transcripts within the same condition. The model jointly captures codon-level occupancy and transcript-level mean ribosome load using a compact architecture, achieving stable and accurate predictions.

## Supporting information

results for GSE133393 are presented in the Supplementary Materials

## Notes

https://github.com/yaozhong/riboPipe

### Competing Interest Statement

The authors have declared no competing interest.

### Summary of Updates

reference inconsistence and incompleteness have been revised

