## Supplementary material for "RiboPipe: efficient per-transcript codon-resolution ribo-seq coverage imputation for low-coverage transcripts": results for GSE133393 are presented in the Supplementary Materials

#### Supplementary Figure S1. Example case studies of ribosome occupancy reconstruction

Representative case studies illustrating predicted and observed ribosome coverage profiles across transcripts in the test set. The predicted codon-resolution ribosome occupancy closely follows the experimentally observed signals, demonstrating that RiboPipe accurately reconstructs both global coverage patterns and local ribosome occupancy peaks.

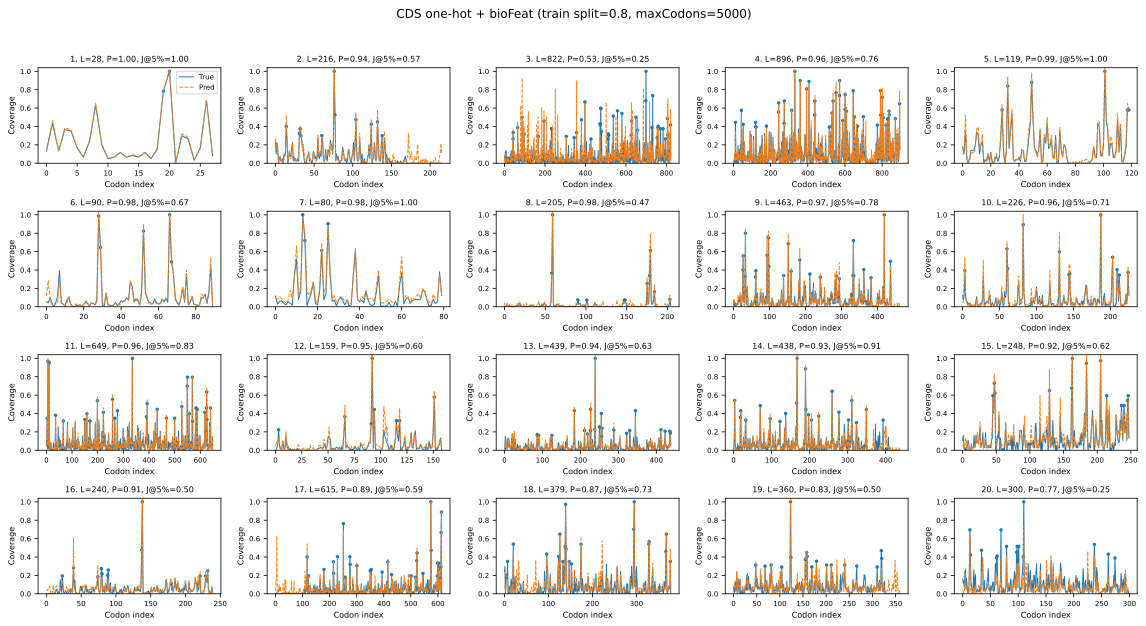

### S1. Analysis on the GSE133393 Dataset

To further validate the performance of RiboPipe on an independent dataset, we applied the framework to the GSE133393 ribosome profiling dataset generated in HEK293 cells. The same preprocessing and evaluation protocol described in the main text was used.

#### S1.1 Data preprocessing and pipeline execution

Raw ribosome footprint reads were aligned to annotated coding sequences derived from the GENCODE v47 transcript reference (gencode.v47.transcripts.fa). P-site assignment was applied to obtain codon-resolution ribosome occupancy profiles. Transcripts were ranked by total CDS ribosome coverage and filtered according to the P75 threshold as described in the main text.

Example pipeline commands:

ribopipe preprocess --csv GSE133393_HEK_Psite_rawcount.csv --fasta gencode.v47.transcripts.fa --out-dir GSE133393_out

ribopipe matrix --npz-dir GSE133393_out --out-csv GSE133393_out/coverage_matrix_transcript_x_sample.csv

ribopipe biofeat --cds-npz GSE133393_out/sample.npz --trna-json trna_copy_numbers.json --out-npz GSE133393_out/bio_features.npz

ribopipe train_pipeline --coverage-csv GSE133393_out/coverage_matrix_transcript_x_sample.csv --npz-dir GSE133393_out --bio-feat GSE133393_out/bio_features.npz --threshold P75 --epochs 200 --train-split 0.8 --max-codons 5000

#### S1.2 Model performance on GSE133393

The RiboPipe model achieved stable and accurate reconstruction of codon-resolution ribosome occupancy profiles on the GSE133393 dataset. Key metrics are summarized below.

Coverage Pearson correlation: 0.823

Coverage Spearman correlation: 0.688

MRL Pearson correlation: 0.863

Binary accuracy: 0.868

Peak Jaccard index (top 5% codons): 0.565

#### S1.3 Case study visualization

Predicted ribosome coverage profiles for representative transcripts are shown in Supplementary Fig. S1. The predicted codon-resolution coverage closely follows the observed ribosome occupancy signals across transcripts.

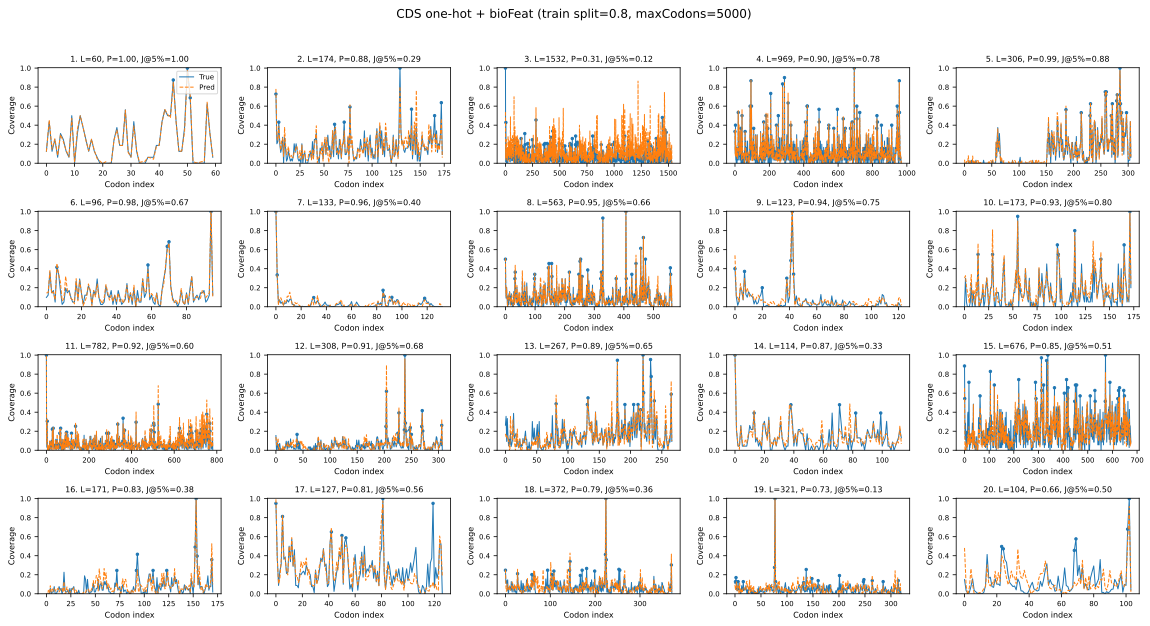

#### S1.4 Ablation analysis on GSE133393

Ablation experiments were performed to evaluate the contribution of biologically informed sequence features (bioFeat), peak-weighted loss (WMSE), and mean ribosome load supervision (MRL).

| Model Variant | Coverage Pearson | Coverage Spearman | MRL Pearson | MRL MSE | BinAcc | Peak Jaccard (Top 5%) |
| --- | --- | --- | --- | --- | --- | --- |
| Default (Full model) | 0.823 | 0.688 | 0.863 | 0.00098 | 0.868 | 0.565 |
| noBioFeat | 0.813 | 0.674 | 0.850 | 0.00112 | 0.862 | 0.539 |
| noWMSE | 0.821 | 0.705 | 0.853 | 0.00110 | 0.855 | 0.537 |
| noMRL | 0.822 | 0.686 | -0.161 | 0.01907 | 0.869 | 0.561 |
| noWMSE + noMRL | 0.825 | 0.711 | -0.403 | 0.01974 | 0.867 | 0.555 |

These ablation results indicate that biologically informed sequence features improve coverage reconstruction, peak-weighted optimization improves peak detection, and joint MRL supervision primarily contributes to transcript-level translation modeling.
